# Biodiversity databases as underutilized resources for pathogen discovery: a quantitative synthesis of bat and rodent tissue collections in natural history museums

**DOI:** 10.1101/2025.11.28.691153

**Authors:** Colleen E. Cronin, Maya M. Juman, Alexander J. Richardson, Christopher A. Whittier, Daniel J. Becker, Adam W. Ferguson

## Abstract

Zoonotic spillovers are becoming increasingly frequent, and the devastating effects of the SARS-CoV-2 pandemic demonstrate our continued inability to combat their consequences effectively. Natural history museums can enhance the study of zoonoses by serving as valuable resources for understanding the ecology and evolutionary history of pathogens and their wildlife hosts. Despite the growing interest in the role museums can play in pathobiology research and zoonotic risk assessment, there remains a lack of centralized resources for locating tissue samples that may be leveraged for pathogen discovery. Using the world’s most significant global aggregator of museum specimens, The Global Biodiversity Information Facility (GBIF), we examined how such tools could be adopted to identify specimens that might be sources of viral genetic material. Focusing on tissue samples from the mammalian orders Rodentia and Chiroptera, speciose taxa that host a high diversity of known zoonotic viruses, we examined temporal, spatial, and taxonomic gaps and patterns in the available tissue samples. Our analyses reveal a heavy bias toward tissue samples collected from the Americas (and consequently, taxonomic groups found in the Americas), with most collected samples housed in North American institutions. This limits the scope of future pathogen discovery efforts and presents a barrier to pandemic preparedness in the Global South. We also examine gaps in metadata quality (e.g., descriptions of preservation method and storage medium) and outline recommendations for GBIF to facilitate future biosurveillance projects and effectively incorporate natural history museums into One Health disease research.

**Author Summary:** The SARS-CoV-2 pandemic has driven a greater research focus on understanding wildlife reservoirs of zoonotic pathogens. As a complement to field-based sampling of wildlife, natural history museum collections house millions of specimens that could be used by researchers to study pathogens quickly, safely, and cost-effectively. Digital databases of museum specimens and their associated tissue samples were originally created for biodiversity research, but these could be adapted to guide pathogen discovery research. Using the largest of these databases, The Global Biodiversity Information Facility, we explored tissue samples from rodents and bats, which have been shown to carry a disproportionate number of zoonotic pathogens. We searched for keywords that would indicate the presence of relevant tissue samples and then mapped the results visually. This revealed disproportionate distributions of tissue samples across time, space, and taxonomy. Our study is the first attempt to assess how a biodiversity database can be used for this novel purpose. We suggest changes that would improve these databases for zoonotic disease research.

## Introduction

Most emerging infectious diseases are zoonotic in origin (1), but the ecology of these pathogens and the exact drivers of their emergence remain elusive (2). This gap in knowledge has resulted in solutions that are often reactive rather than preventative, which limits our ability to combat rapidly emerging zoonoses (3). The first step in predicting and eventually mitigating spillover is identifying existing and likely host species.

Wild animals harbor thousands of viruses that have yet to be discovered, many of which likely have zoonotic potential (2). Species from the orders Rodentia and Chiroptera have the highest number of reported zoonotic viruses; this is at least in part a function of the high species richness in both orders (5), although some studies suggest that even after controlling for sampling bias and species richness, bats have a significantly higher proportion of zoonotic viruses than all other mammalian orders (6, 7). As common hosts for zoonotic viruses, bats and rodents are important groups for exploring limitations or gaps in resources for studying zoonotic pathogens. Within these orders, certain species are more likely to carry and transmit zoonotic viruses. Studies have linked zoonotic pathogen richness with larger ranges and more litters at a younger age in rodents (8) and range overlap, smaller litters, more litters per year, and longevity in bats (9, 10). This suggests that species traits can be used as a guide to predict zoonotic risk and focus research efforts on higher-risk mammalian species.

Host–virus associations are crucial data that allow researchers to better predict and mitigate spillover risk, such as the spatial and host range of a pathogen (11, 12, 13). Understanding the ecological and evolutionary relationship between a host and virus can also provide valuable insights into cellular interactions, changes in prevalence and distribution, and factors that influence the likelihood of spillover into humans (14, 15). However, the rarity of many viral infections and high spatiotemporal variation in prevalence necessitate the collection and screening of large sample sizes from hosts (16), which can involve prohibitively expensive and time-consuming field work.

In contrast to prospective field sampling, natural history museums already hold an estimated 1.1 billion specimens around the globe (17). Mammal collections in North America alone contain more than five million specimens (18), and new specimens are added each year (19). In the last decade, there has been growing interest in using these specimens for pathobiology research. Natural history museums are thereby uniquely positioned to circumvent many of the issues involved in field-based sampling, and they also enable researchers to cheaply and efficiently study pathogen occurrence and diversity across space and time (19, 20, 21, 22, 23, 24).

A classic example of collections-based viral host identification is the Sin Nombre hantavirus that emerged in 1993 in the American Southwest and resulted in 27 human fatalities. Researchers used specimens from the Museum of Southwestern Biology to not only identify the definitive reservoir as the common deer mouse (*Peromyscus maniculatus*) but also determine that the virus had been circulating in wild deer mouse populations for at least a decade prior to the first known spillover event (25, 26). Since then, over 40 hantaviruses have been discovered in various small mammals, many with the use of museum specimens, and archived tissue samples have provided new insights into the evolutionary origins, genetic diversity, and spatial and temporal distribution of these viruses (20, 27). In more recent studies, museum specimens have been used to study RNA viruses (28, 29), DNA viruses (30), bacteria (31), and metazoan parasites (32). Recent methodological advances have even facilitated the recovery of RNA from formalin-fixed specimens (33), an approach that could enable viral screening of much older material. However, despite mounting evidence that museum collections are critical resources for the identification, tracking, and mitigation of zoonotic pathogens, they remain largely overlooked by disease ecologists and virologists (19).

Incorporating natural history museums into One Health research also presents its own set of challenges. Most specimens were collected at times when the research questions being asked today were unimaginable (16). Traditional specimen preservation methods (e.g., formalin fixation; storage in ethanol, DMSO, or EDTA) are insufficient for preventing the degradation of viral genetic material, particularly RNA. In addition to long-term storage requirements, the utility of a tissue sample for viral research depends on the initial storage methods in the field as well as any changes during transport (34). Ideally, samples would be flash-frozen in liquid nitrogen as soon as possible (35), either without a buffer (e.g., if collecting tissue for host genomics) or with a buffer that preserves genetic material but inactivates live virus (e.g., DNA/RNA Shield) (36). After initial storage, freeze–thaw cycles should be avoided (34). However, prior to 2014, there were no best-practice standards for the documentation and storage of genetic resources in natural history museums, and storage methods varied greatly across collections (34). Moreover, the Systematic Collections Committee of the American Society of Mammalogists only formally published their standards and guidelines on genetic resource maintenance in 2019 (35).

Despite this lack of guidance, museums have made great strides in recent decades to embrace the concepts of the “holistic specimen” (37), “extended specimen” (38), and “open specimen” (39). As a result, the preservation of tissue samples, metazoan parasites, and other associated microbiological samples is becoming increasingly common (24, 37). However, for these samples to be useful to the broader scientific community, the relevant metadata must also be archived in a publicly accessible digital format, and the push to digitize and share these data lags behind that of primary biodiversity metadata (40). For many decades, museums held their digital archives on various internal databases, each using different recording formats and software packages (37, 40). The standardization and open access of these data remain pressing challenges for natural history museums in the context of One Health (37, 41).

Publicly available biodiversity databases address many of these issues (42), and each database often has a specific focus related to spatial coverage and record type (40). Early efforts to digitize and mobilize museum data were also often taxon-specific, such as the Mammal Networked Information System (43). Today, the Global Biodiversity Information Facility (GBIF) is one of the largest biodiversity databases in the world, with over one billion records from institutions in over 50 countries (40, 44). By implementing standard terminology such as Darwin Core (45) and new tools such as the Integrated Publishing Toolkit (46), GBIF has allowed museums to easily share their data online (37). Most importantly, GBIF works as a downstream data aggregator, collecting data from a network of biodiversity databases, organizations, and data papers such that information can be shared and searched in a centralized fashion (40). Many researchers have heralded the potential of data aggregators such as GBIF to facilitate use of museum collections in other research initiatives (19, 23, 37, 41, 44, 47). However, studies that quantitatively evaluate the spatial, temporal, and taxonomic coverage of GBIF for the purpose of pathobiology are lacking.

Here, we test whether biodiversity data aggregators such as GBIF can be adopted for users to identify suitable tissues for pathobiological studies. We examine temporal, geographic, and taxonomic representation in digitized bat and rodent tissue samples that could be used for pathogen surveillance, including those from species predicted to carry the most undescribed zoonotic viruses (6). Finally, we outline recommendations for museums and database aggregators to improve data accuracy and facilitate the inclusion of natural history museums in One Health research.

## Results

Of the 140 museum collections or institutions with tissue samples in GBIF, 26 showed evidence of possessing only bat tissue samples, 32 showed evidence of possessing only rodent tissue samples, and 89 showed evidence of possessing both bat and rodent tissue samples. In total, 588,425 (out of 4,586,821; 12.8%) listed specimens had associated tissue samples, of which 475,834 were from rodents and 112,591 were from bats (see Supplementary Data S1 for the full list of institutions and their associated tissue samples).

Among specimens with associated tissue samples, the search term “tiss” or “tejid” yielded the highest number of occurrences, appearing in over 59% of the dataset. Over 26% of tissue samples appeared to be in ethanol, while ∼45% were frozen. Of reported tissue types, liver samples were the most common, followed by kidney, heart, spleen, lung, muscle, and colon (Table 1).

**Table 1.** Occurrence of preservation method and tissue type in total dataset and breakdown by taxonomic order.

| Term | Total occurrences | Rodent occurrences | Bat occurrences |
| --- | --- | --- | --- |
| “blood” / “sangre” / “serum” / “suero” / “plasma” | 66,382 | 64,995 | 1,387 |
| “tiss” / “tejid” | 348,165 | 275,050 | 73,115 |
| “biopsy” / “biopsia” | 1,169 | 7 | 1,162 |
| “swab” / “torunda” | 13,778 | 13,178 | 600 |
| “ethanol” / “etanol” / “EtOH” | 157,748 | 125,099 | 32,649 |
| “froze” / “freez” / “congelad” | 266,471 | 237,675 | 28,796 |
| “DMSO” / “EDTA” / “VTM” / “RNAlater” / “shield” / “buffer” / “lysis” | 2,217 | 1,531 | 686 |
| “heart” | 102,393 | 92,147 | 10,246 |
| “kidney” | 108,239 | 97,394 | 10,845 |
| “liver” | 150,432 | 136,200 | 14,232 |
| “lung” | 67,402 | 64,124 | 3,278 |
| “muscle” | 30,383 | 24,004 | 6,379 |
| “spleen” | 72,708 | 69,345 | 3,363 |
| “colon” | 1,010 | 718 | 292 |

### Temporal distribution

The collection year of all bat and rodent samples in GBIF (not just those with associated tissue samples) ranged from 1591 to 2025, with a median year of 1972 that corresponded with an uptick in collecting (Fig. 1). Overall, 391,809 samples (8.5%) were missing collection date information. For samples with associated tissues, the years of collection ranged from 1806 to 2025, with a median collection year of 2005. Of these samples, 15,108 (2.6%) were missing collection date information.

**Fig. 1.**
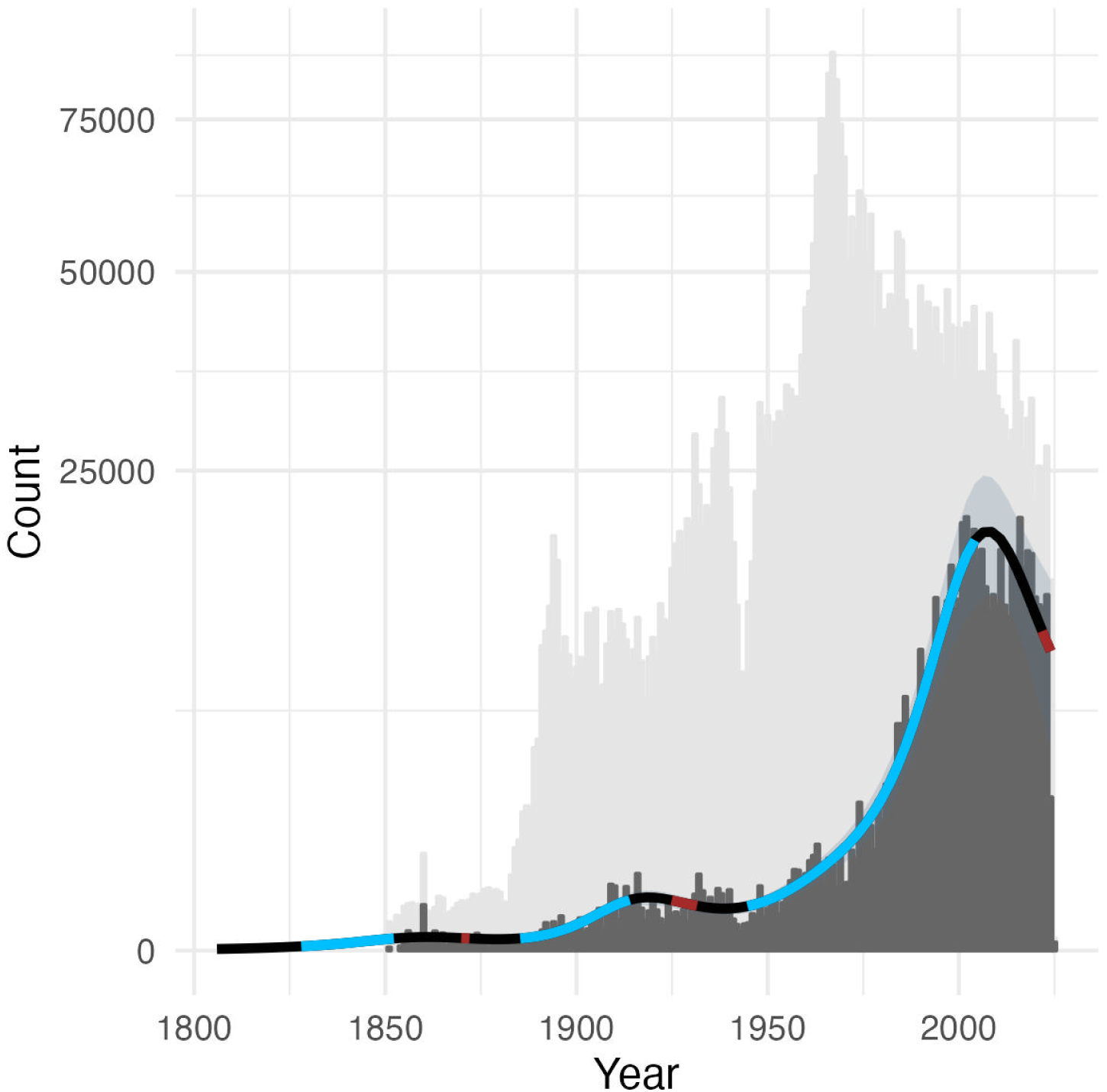
Barplot indicating the collection year of all bat and rodent specimens in GBIF (light grey) and those with associated tissue samples (dark grey), with the number of samples per year shown on a square root scale. The black line indicates the fitted spline from a GAM of tissue samples collected per year. Blue intervals indicate periods of significant increase in tissue collecting while red intervals indicate periods of significant decline.

We fitted a generalized additive model (GAM) to the tissue samples collected per year to identify periods of significant increase or decrease in the rates of tissue collection up until 2024 (48). From 1828 to 1852, tissue collection showed a significant increase, followed by a brief decline in the late 1860s. Tissue collection increased again from 1885 to 1914, followed by another brief decline from 1925 to 1932. After 1945, tissue collections grew rapidly until 2004, but since 2021 the rate of tissue collecting has been declining.

### Spatial biases

The dataset included tissue samples from 190 countries or territories with unique country codes. An additional 376 samples were from unknown countries (i.e., country code ZZ). However, sampling effort was highly heterogeneous across countries/territories, ranging from one sample (i.e., Netherlands Antilles, Bahrain, Bosnia and Herzegovina, Belarus, Cyprus, Georgia, Guam, Maldives, Oman, Saint Helena, Ascension and Tristan da Cunha, Slovenia, and Tajikistan) to 349,890 (i.e., the USA) (Fig. 2). The second most sampled country was Mexico, with 22,830 tissue samples. A generalized linear model (GLM) of binary sampling effort across all regions (i.e., Africa, Americas, Asia, Europe, Oceania) revealed disproportionate sampling of African, American, and Asian countries (*χ*^2^ = 14.58, *p* = 0.0057, *R*^2^ = 0.05) (Supplementary Fig. 1). The number of samples per country also varied significantly across subregions (*χ*^2^ = 2,214,834, *p* < 0.0001, *R*^2^ = 0.62), which was driven by the disproportionate number of samples from the USA, Mexico, and Canada (Supplementary Fig. 2).

**Fig. 2.**
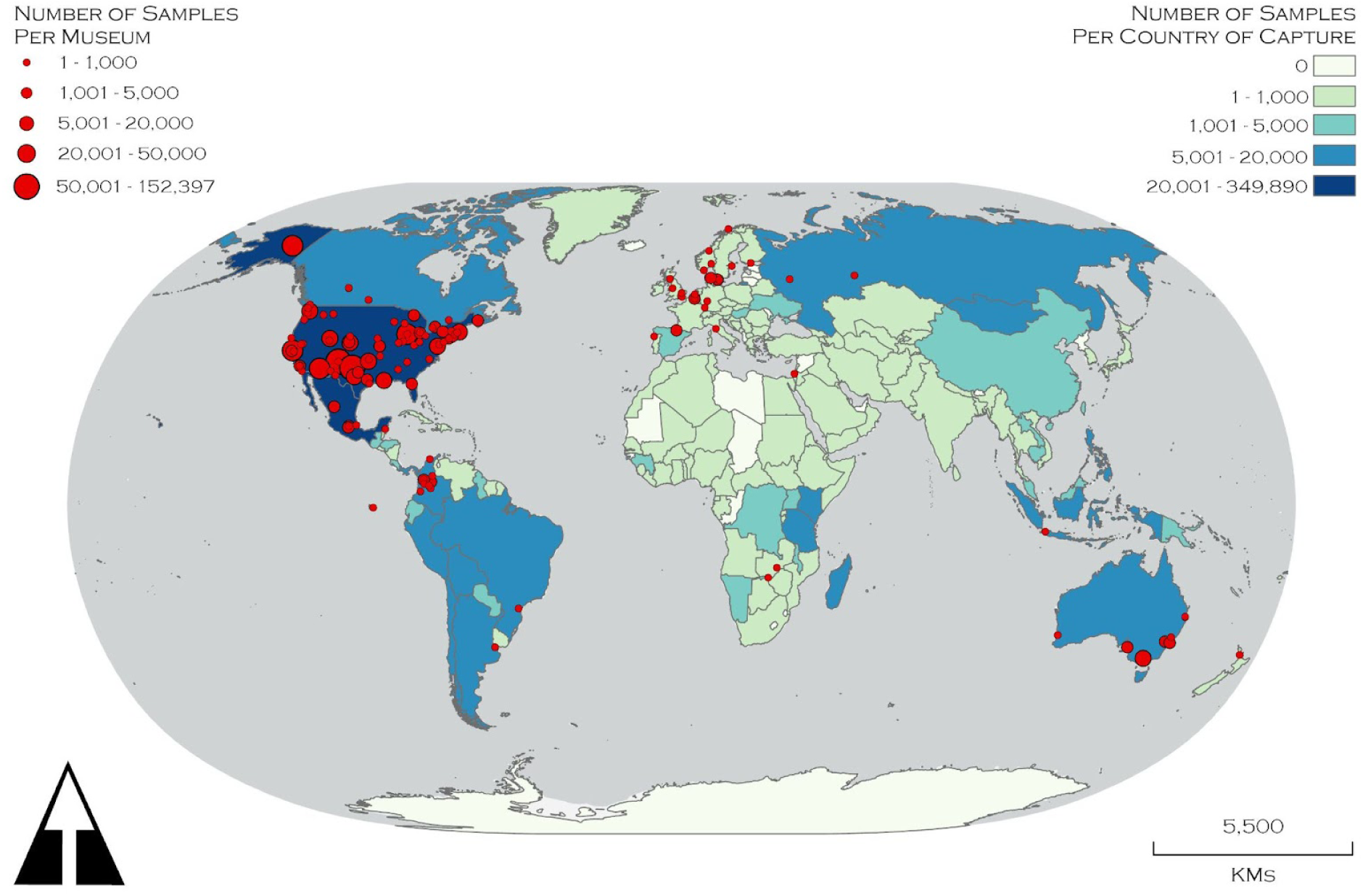
Global map showing the spatial distribution of bat and rodent specimens with associated tissue samples in GBIF as of July 14, 2025. Countries of collection are shown with shading, while red dots indicate the locations of museum collections in the dataset. The map was constructed in ArcGIS Pro (49) with the World Natural Earth II projection (50) and the World Light Gray Base basemap (51). ISO two-digit country codes were used to match our dataset to the World Countries (Generalized) feature layer provided by Esri via ArcGIS Living Atlas of the World (52).

Our analysis also revealed strong biases in the spatial distribution of reported tissue holdings (Fig. 2). The size of museum collections was heterogeneous across the 26 countries represented in the dataset, ranging from two samples (i.e., France, the Netherlands) to 533,322 samples in the USA. A GLM across all regions revealed disproportionately high fractions of countries with tissue holdings in Europe and the Americas (*χ*^2^ = 20.10, *p* < 0.001, *R*^2^ = 0.11), and the number of samples per country also varied significantly across subregions (*χ*^2^ = 2,478,857, *p* < 0.0001, *R*^2^ = 0.77), driven by a disproportionate number of samples in North American museum collections (Supplementary Figs. 3,4).

### Taxonomic biases

The dataset initially listed 2,445 species and 619 genera of bats and rodents with associated tissue samples. Of these, 290 did not match to our taxonomic backbone (53). After manual taxonomic reconciliation using the Mammal Diversity Database, 47 of these 290 taxa were recently described or otherwise not in the mammalian phylogeny and could not be harmonized. This left us with 2,377 harmonized species with tissue samples, including 842 bat species and 1,416 rodent species. Based on our taxonomic backbone (53), these samples represent approximately 65% (842/1,287) and 59% (1,416/2,392) of recognized bat and rodent species, respectively.

We found intermediate phylogenetic signal in binary sampling effort across bat species (*D* = 0.90), departing from both phylogenetic randomness (*p* < 0.001) and Brownian motion models of evolution (*p* < 0.001). We then applied phylogenetic factorization, a graph-partitioning algorithm, which revealed that the superfamily Noctilionoidea (containing the families Myzopodidae, Mystacinidae, Thyropteridae, Noctilionidae, Furipteridae, Mormoopidae, and Phyllostomidae) was overrepresented, while the genera *Scoteanax*, *Harpiola*, *Murina*, and *Scotozous*—a subclade of Old World vesper bats—were underrepresented (Fig. 3a; Table 2a). Among sampled species, the number of tissue samples per bat species ranged from one to 4,726 and was distributed across the bat phylogeny with intermediate phylogenetic signal (Pagel’s λ = 0.28), departing from both Brownian motion models of evolution (*p* < 0.001) and phylogenetic randomness (*p* < 0.001). Eleven *Myotis* species—all North American members of the subgenus *Pizonyx*—were overrepresented relative to other taxa; however, the genera *Artibeus* and *Dermanura* were underrepresented (Fig. 3b; Table 2b).

**Fig. 3.**
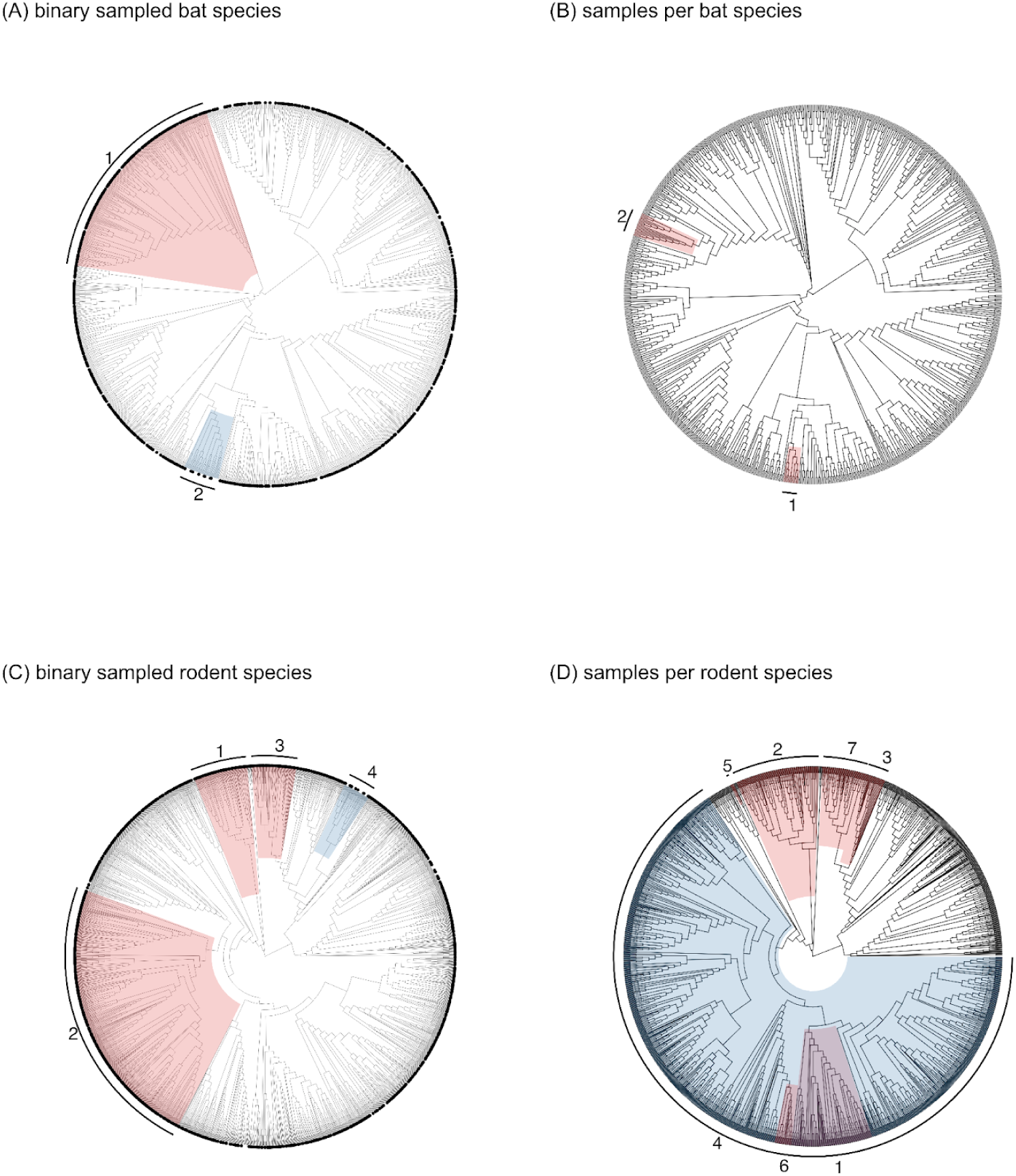
Host and viral taxonomic biases in pteropodid–paramyxovirus sampling effort. (A) Whether or not a bat species has tissue samples in our dataset, across all Chiroptera (black dots indicate species that have been sampled); (B) logged number of tissue samples per sampled bat species; (C) whether or not a rodent species has tissue samples in our dataset, across all Rodentia (black dots indicate viruses that have been detected); (D) logged number of tissue samples per sampled rodent species. Red and blue shading indicates clades with greater or lesser sampling effort, respectively, in comparison to all other taxa (as identified by phylogenetic factorization), with clade numbers corresponding to Table 2.

**Table 2.**
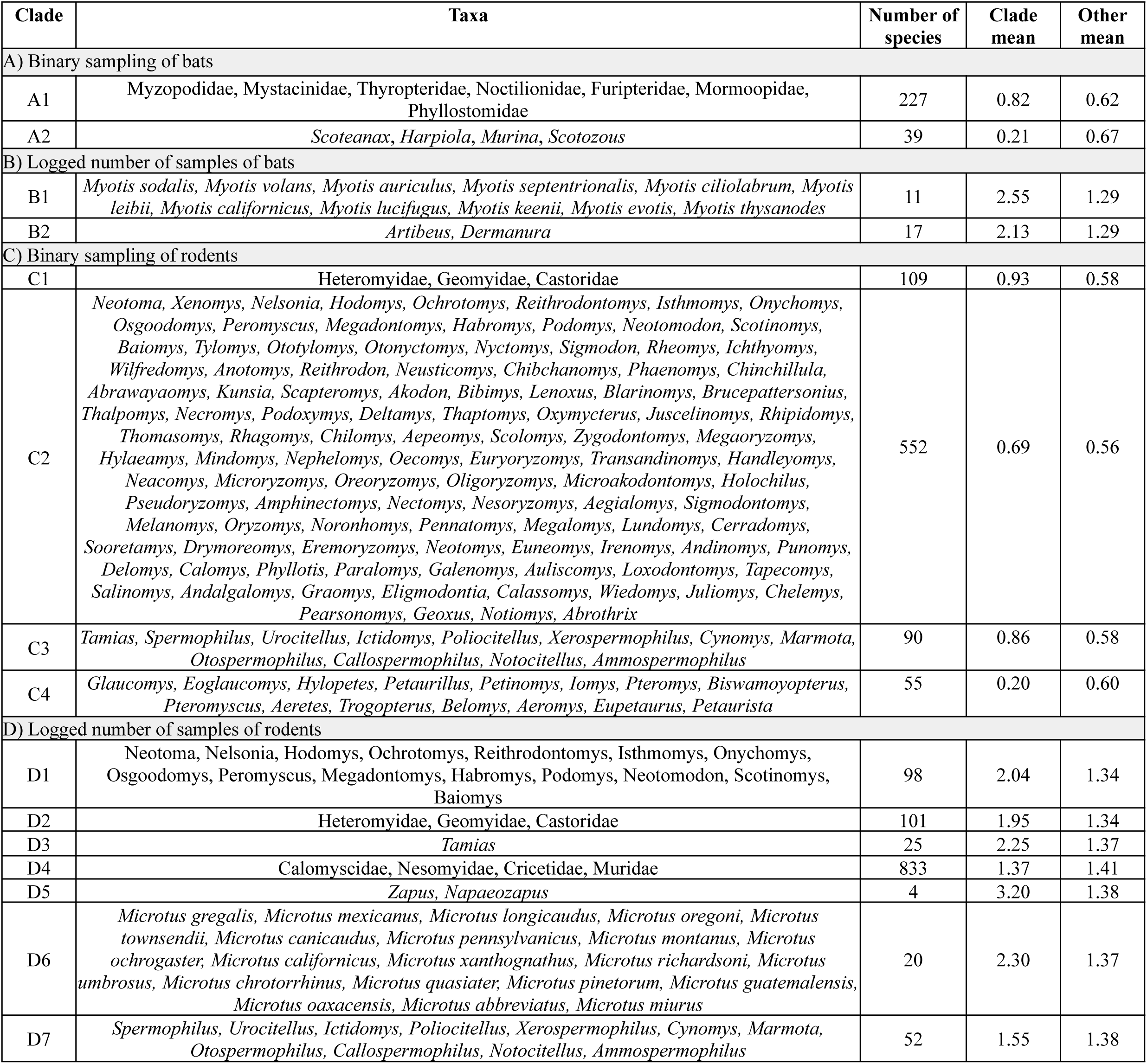
Bat and rodent clades that differ significantly from the rest of their respective trees in terms of different measures of sampling effort, as identified by phylogenetic factorization. (A) Whether or not a bat species has tissue samples in our dataset, across all Chiroptera (0: not sampled; 1: sampled); (B) number of tissue samples per sampled bat species; (C) whether or not a rodent species has tissue samples in our dataset, across all Rodentia (0: not sampled; 1: sampled); (D) number of tissue samples per sampled rodent species (Fig. 3). We report all significant clades detected through phylogenetic factorization (identified with Holm’s sequentially rejective test at a family-wise error rate of 5%), the taxa and number of species within those clades, and the mean sampling effort for the clade compared to the rest of the tree. Bat and rodent taxonomy follows the most recent mammal phylogeny (53).

Similarly, we found intermediate phylogenetic signal in binary tissue sampling effort of rodents (*D* = 0.82), departing from both phylogenetic randomness (*p* < 0.001) and Brownian motion models of evolution (*p* < 0.001). Phylogenetic factorization revealed that some rodent clades (including the largely North American families Heteromyidae, Geomyidae, and Castoridae in the suborder Castorimorpha, as well as some genera in Cricetidae) have higher fractions of sampled species relative to other clades, while some genera have disproportionately lower fractions of sampled species (Fig. 3c; Table 2c). Among sampled species, the number of tissue samples per rodent species ranged from one to 54,687 and was distributed across the phylogeny with intermediate phylogenetic signal (Pagel’s λ = 0.49), departing from both Brownian motion models of evolution (*p* < 0.001) and phylogenetic randomness (*p* < 0.001). For example, rodents in the families Heteromyidae, Geomyidae, and Castoridae (members of suborder Castorimorpha) were overrepresented in terms of sampling intensity relative to other taxa, while those in Calomyscidae, Nesomyidae, Cricetidae, and Muridae (clade Eumuroida) were underrepresented (Fig. 3d; Table 2d).

In 2017, Olival et al. used species-level trait data to predict the risk of spillover from other mammals into humans (6). Our dataset included tissue samples for the ten rodent and ten bat species with the highest predicted number of undiscovered zoonotic viruses (6), suggesting that GBIF can be used to locate samples for testing predictions (Table 3).

**Table 3:**
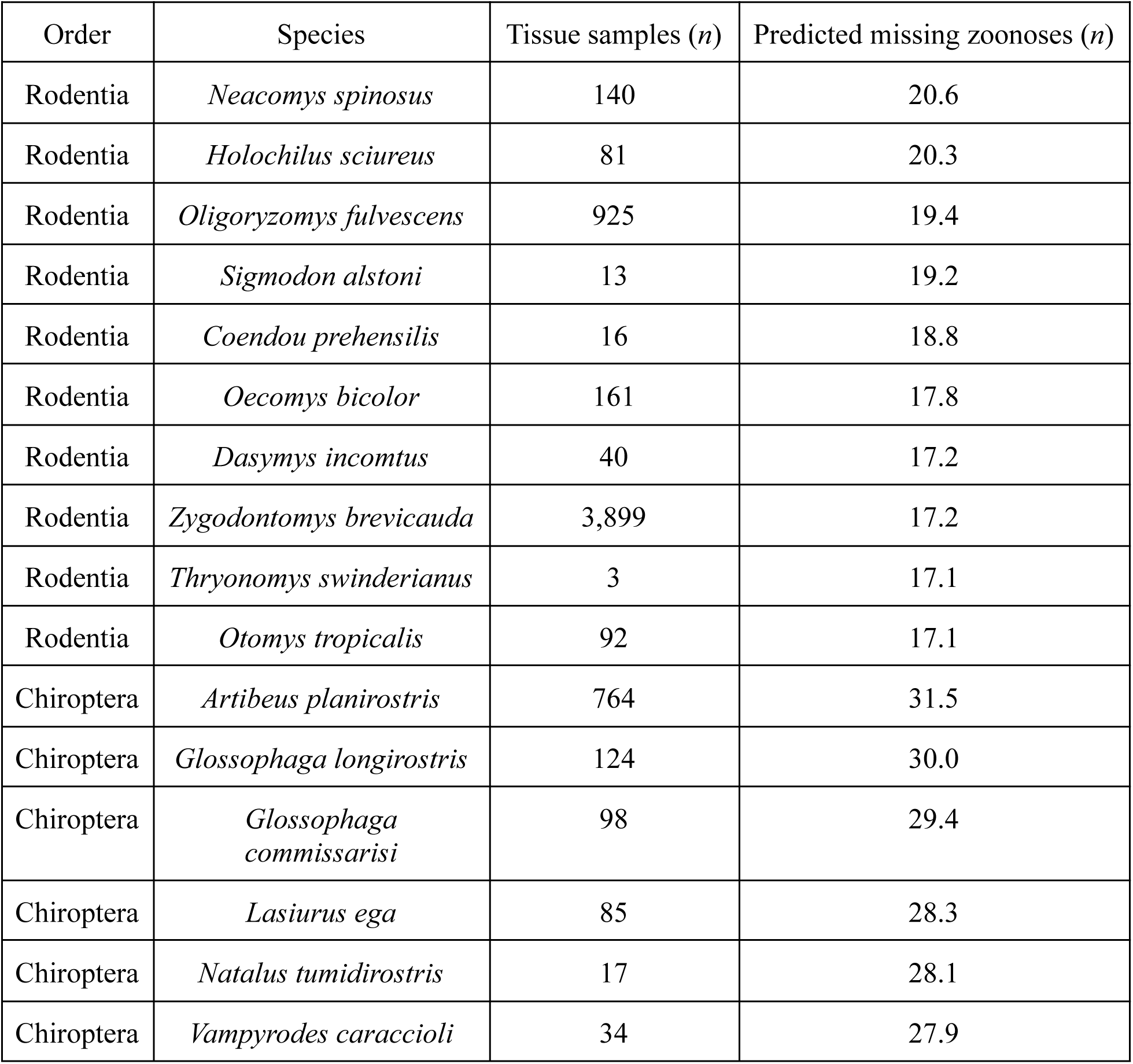
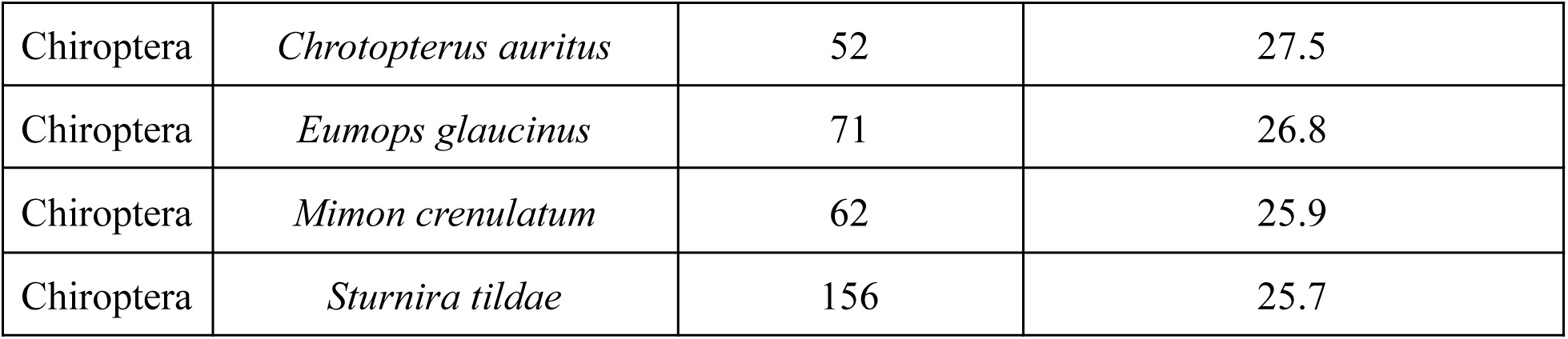
Potentially viable tissue samples from the ten bat and ten rodent species with the highest number of predicted missing zoonoses as modeled by Olival et al., 2017 (6). A full list of museums housing these samples is presented in Supplementary Data S2.

## Discussion

Biodiversity databases have the potential to guide pathogen discovery and surveillance efforts, as they consolidate metadata on archived samples from wildlife globally. We used one of the most comprehensive of these databases, GBIF, to evaluate temporal, spatial, and taxonomic gaps in rodent and bat tissue holdings at natural history museum collections and other biorepositories reported in GBIF. Of the roughly 4.5 million rodent and bat records in GBIF, less than 13% (*n* = 588,425) included a term indicating a preserved tissue, serum, or biopsy sample that could potentially be used for pathogen diagnostics. Of that subset of records, only around half appeared to be frozen (Table 1).

Tissue samples were collected from 1806 onward, with a median collection date of 2005 (Fig. 1). This coincides with a shift toward the “extended specimen” model of collections, where associated tissues are preserved along with a voucher specimen, often in freezers or liquid nitrogen (54). The paucity of tissue samples from the 19th and first half of the 20th century limit our ability to use these collections as “time machines” for viral diagnostics further back than a century, although other pathogens like metazoan parasites are detectable (32). The continued collection and mindful preservation and storage of new biological materials represents an investment in future pathogen detection and disease prevention efforts. However, the growth of bat and rodent tissue collections has slowed in recent years (Fig. 1), threatening future pandemic preparedness and hindering our ability to study the effects of ongoing global change (20, 37, 55, 56, 57). Despite the obvious value of collections, the costs needed to collect and maintain voucher specimens have made natural history museums vulnerable to changes in funding and resource prioritization (37, 58, 59, 60, 61, 62, 63, 64, 65). Collections at academic institutions have been especially impacted as a result of state budget cuts and shifting interest to other scientific branches (66). Further, an increased focus on nonlethal samples or observations (e.g., camera-trap data, fur/skin samples, etc.) has contributed to the decline in collection rates, but these approaches are less valuable for pathogen discovery efforts than whole specimen collection (55, 57).

To effectively use GBIF and similar databases as resources for pathogen surveillance, researchers must be able to locate appropriately preserved specimens for DNA and RNA extraction (or other methods such as serology). The lack of standardized reporting of preservation and storage details in GBIF is a serious barrier to selecting target specimens and tissues for data collection. Of the reported tissue samples, we found that over one-quarter are preserved in ethanol. If these fluid-preserved specimens were originally fixed in formalin, this presents an issue for nucleic acid recovery (particularly RNA, which is problematic for the detection of pandemic-causing RNA viruses such as coronaviruses and paramyxoviruses) (29). New methodological advances are enabling us to extract nucleic acid material from older, formalin-fixed specimens (33, 67, 68). However, this work is early in development and is more expensive than traditional RNA extraction; therefore, the inclusion of more details in GBIF about preservation methods would help facilitate these preliminary projects.

The tissue samples in our dataset were heterogeneously distributed across space, in terms of both country of origin and institution country. Countries in the Americas were overrepresented from multiple perspectives (Fig. 2). This bias has potential downstream impacts on our future ability to screen tissues from a broad range of regions for potential emerging pathogens. Europe, Asia, and Africa are all predicted hotspots for emerging zoonotic spillover events (1), but the lack of samples with associated tissues from these continents may hinder the sample size of future museum-based surveillance studies.

Such results are consistent with the current literature on where viral research related to bats in particular is lacking. Western Asia is home to 96 bat species, yet a recent study found no publications investigating bat-associated coronaviruses in most of the countries there (69). Despite the high diversity of bat species in Central Africa and Southeast Asia, countries in these regions are also severely underrepresented in the bat literature (69). Similarly, a review of global coronavirus sampling in rodents and other small mammals found striking sampling gaps in Western and South Asia, Eastern Europe, Australia, and much of the African continent (70). Our results further support the observation that tissue samples of bat and rodent specimens are largely deficient in these regions.

There is also a dramatic skew in the locations of museum collections with bat and rodent tissue holdings, with most samples housed in North American institutions (Fig. 2). The concentration of these resources in the Global North limits collection-based pathogen surveillance and discovery efforts in underrepresented regions. Building biorepository capacity in the Global South will be critical in fostering a more collaborative, fair, and comprehensive approach toward pathogen spillover prediction and management (71).

Bat and rodent tissue samples in GBIF were also skewed in their taxonomic distributions. Bats within the superfamily Noctilionoidea show higher proportions of sampled species relative to other clades. Most of the families within this clade are found only in the Americas (e.g., including but not limited to Phyllostomidae and Mormoopidae), which is consistent with the overrepresentation of North and South American countries in the dataset. However, many Asian, African, and Oceanian bats are the demonstrated or hypothesized reservoirs of zoonotic viruses like coronaviruses (72), paramyxoviruses (73), and filoviruses (74), suggesting that this sampling bias might limit effective surveillance across all bat taxa for potential epidemic-causing viruses. We identified a similar taxonomic and geographic skew in rodents, where species in the largely New World families Heteromyidae, Geomyidae, and Castoridae are sampled more frequently than those in other clades.

Predicting pathogen spillovers is a major research topic, and recent studies have employed predictive models that leverage host–pathogen association data for this purpose (6, 9, 11, 75). The tissue samples in GBIF can be used to test such predictions. For example, there are 764 and 140 samples, respectively, from *Artibeus planirostris* and *Neacomys spinosus*, the bat and rodent species predicted to carry the most undetected zoonotic viruses (6). By screening tissue samples from these specimens, researchers could empirically test such predictions, following a model-guided viral discovery framework (29). In turn, novel detections could be iteratively integrated back into such models to improve forecasts and potentially mitigate the risk of zoonotic spillover before it occurs (14, 75).

Given the availability of such tissue samples, the data from our study can be used as a complementary approach to fieldwork, which would augment zoonotic pathogen research by expediting the timeline and reducing the cost of studying emerging infectious disease risks. Our results can also guide prospective collection efforts by illuminating which species and regions have already been extensively sampled and which are lacking representation in museum collections. Prioritizing subsequent collection efforts in areas that are species-rich but underrepresented would greatly facilitate future pathogen discovery projects.

Our study is not without limitations. Although GBIF is an international database with information from institutions in over 50 countries, our results were limited to one database aggregator. As publishing data in GBIF is voluntary, many museums with viable tissue samples were potentially overlooked, especially smaller collections which may not be participating members of GBIF. Specimens collected by smaller museums or research teams are less likely to be properly recorded, and these “dark data” often remain hidden from the larger research community (76). Of the collections that have been digitized, many are not integrated into the network of databases or have inaccurate or outdated information (41, 76). Even records that have been incorporated into major databases shared with GBIF are often missing vital data, including but not limited to the date or location of collection (44). Furthermore, these databases and data aggregators were designed for the purpose of biodiversity research (47). As a result, standards regarding the inclusion and entry of data that are critical to searching for appropriate samples for pathobiology research (e.g., preservation method or buffer type) are commonly lacking.

Within GBIF, it is also possible that suitable samples could have been missed in our text mining search. Lack of data, including country name, tissue presence, and preservation details, would have led to omission of potentially relevant results. Inconsistent record keeping and lack of standardization also limited the feasibility of a fully exhaustive study. All researchers on our team were native English speakers and, despite our efforts to include the most common languages present in the dataset, this bias could have influenced the search terms we selected. Finally, we caution that it is not possible to extrapolate enough details about tissue preservation techniques to say for certain whether these samples are all suitable for pathobiology research (e.g., lack of information surrounding sample preparation after capture and long-term storage, such as temperature, number of freeze/thaw events, and types of buffers used, if any).

These limitations underscore an urgent need for changes in how tissue samples are preserved and how this information is shared across the research community. Researchers have long recognized the need for museums and their associated databases to be integrated with pathogen discovery and research (19, 22, 77). Yet equally crucial is the documentation of these resources to be housed in a centralized portal (78, 79). Large amounts of data, such as those surrounding details of specimen collection and storage, are only useful if they are archived in a standardized and easily accessible manner. While GBIF has made important strides in this direction, it has largely prioritized conservation and biodiversity research, and this focus is reflected in which data are collected and how they are stored. By including the needs of pathogen-oriented researchers in its design, GBIF could further expand its scope to help prevent and mitigate the spillover of zoonotic pathogens. Improved standardization and recording practices, along with a universal portal, will allow researchers to more easily distinguish between actual biases in specimen collections versus gaps in documentation. Proper data storage and sharing will expose which specimens are lacking and which species need to be prioritized. However, separate steps are needed to ensure that, once collected, specimens are used to their greatest research potential.

While the benefits of preserving host voucher specimens and tissue collections have been acknowledged by many within the infectious disease research community, large-scale support has been slow (22). Archives of RNA and/or DNA sequences are now required by most top journals in ecology, evolution, behavior, microbiology, and systematics, whereas specimen deposition is required by only 13% of journals (39). Not only does specimen deposition facilitate repeatability—a foundational tenet of scientific research—but it also allows the specimen to be used in future unrelated studies, particularly if it is properly preserved and stored. Museums serve a larger mission as data stewards and educators of the general public. Incorporating them into the One Health field may catalyze a shift in focus back to this common goal and foster a culture of collaboration as we work to prevent future pandemics.

## Materials and Methods

To quantify the temporal, spatial, and taxonomic distribution of specimens viable for pathobiology research, we queried GBIF on July 14, 2025 for all records in the orders Chiroptera and Rodentia. We filtered data to those records in which occurrence status was listed as present and where the record was categorized as a material sample or preserved specimen (80, 81). We used R 4.2.1 (82) to clean the dataset, including 1) ensuring case uniformity; 2) assigning ISO country codes to avoid discrepancies in country names; and 3) matching codes to generalized country coordinates using a publicly available dataset (83). Several packages, especially those in or peripheral to the *tidyverse* collection (84), were used to import, manipulate, and visualize data, including *dplyr* (85), *tidyr* (86), *ggplot2* (87), *patchwork* (88), and *data.table* (89).

To use the maximum number of data points available for spatial analysis, we focused primarily on country-level capture locations. Rather than exclude records missing exact specimen coordinates, generalized country geolocations were added to records without coordinates. Although we acknowledge this limits visualizing exact within-country collection locations, this approach allows more specimens to be collectively mapped. Of the records with languages listed (2,207,368), English (1,873,092 or 84.9%) and Spanish (278,913 or 12.6%) were the two most common, amounting to over 97% of these records. Therefore, we used these two languages for our keyword searches. To ensure we captured any potential specimens of interest for pathobiology research, we included truncated word forms to account for inconsistency and human error in data entries. We designed our search terms to capture tissue availability and thus included different organs, buffers, and preservation methods.

Using the *stringr* package (90), we searched for tissues in the fields “preparations,” “dynamicProperties,” “occurenceRemarks”, and “materialEntityRemarks” using the following search terms: “blood”, “sangre”, “tiss”, “tejid”, “serum”, “suero”, “plasma”, “biopsy”, “biopsia”, “swab”, “torunda”, “heart”, “kidney”, “liver”, “lung”, “muscle”, “spleen”, “colon”, “ethanol”, “etanol”, “EtOH”, “VTM”, “RNAlater”, “shield”, “lysis”, “DMSO”, “EDTA”, “buffer”, “froze”, “freez”, or “congelad”. We used the results of this search to generate a binary variable indicating the presence or absence of at least one of these search terms in at least one target field.

To map the museums that house these tissues, we used the Google Maps API. Any missing locations after the API query were obtained manually through the GBIF Registry of Scientific Collections, with preference given to the field “institution code.” Duplicates were assumed to be due to acquisitions and manual errors during data entry and were thus combined. We used ArcGIS Pro to visualize the spatial distribution of where bat and rodent specimens with associated tissue samples were collected (49).

To examine temporal patterns in tissue collections, we used the *mgcv* package and restricted maximum likelihood to fit a GAM to the number of tissue samples collected per year (up until 2024). We used penalized thin-plate regression splines to fit year as a nonlinear smoothing term, with number of tissue samples modeled as a negative binomial distribution (91). We then identified periods of significant increase or decline in tissue collection, defined as when the first derivative of the predicted fitted value was significantly nonzero (based on a 95% confidence interval) (48).

To further examine spatial patterns in tissue collection, we fitted two GLMs with collection region (i.e., Africa, Americas, Asia, Europe, Oceania) or subregion (i.e., Australia and New Zealand, Central Asia, Eastern Asia, Eastern Europe, Latin America and the Caribbean, Melanesia, Micronesia, Northern Africa, Northern America, Northern Europe, Polynesia, Southeastern Asia, Southern Asia, Southern Europe, Sub-Saharan Africa, Western Asia, Western Europe) as a categorical predictor. One GLM modeled binary sampling status (i.e., whether or not tissue samples have been collected from a country) as a binomial response, with all countries included and region as a predictor, whereas the other GLM modeled the number of tissue samples per country as a Poisson-distributed response, with only sampled countries included and subregion as a predictor. We also fitted two additional GLMs to assess biases in the spatial distribution of museum holdings. Similarly, one GLM modeled binary holding status (i.e., whether or not a country has any tissue samples) as a binomial response, with all countries included and region as a categorical predictor, whereas the other GLM modeled the number of tissue samples per country as a Poisson-distributed response, with all countries included and subregion as a categorical predictor. We assessed model output with the *car* package (92) and model fit with McFadden’s R^2^ using the *performance* package (93) and visualized differences across regions with the *visreg* package (94).

To assess taxonomic sampling biases in tissue collections, we aligned all sampled bat and rodent species to a mammalian phylogeny (53). This involved relabeling species names to fit this taxonomic backbone where necessary. We first harmonized species names with the *taxize* (95) and *rentrez* (96) packages, followed by subsequent manual relabeling of any remaining taxonomic discrepancies using synonyms listed on the Mammal Diversity Database v2.3 (97). Recently described species missing from the mammal phylogeny were removed for the taxonomic analysis (53).

We then created a binary response variable for all bat and rodent species in the phylogeny, representing whether or not each species is present in our dataset with associated tissue samples. For those that were present, we calculated the total number of samples with tissues associated with each bat and rodent species. To assess phylogenetic signal in sampling effort within each of bats and rodents, we used the *cape*r package (98) to calculate *D* (0: phylogenetic clustering under Brownian motion; 0-1: intermediate phylogenetic signal; 1: phylogenetically random sampling distribution; >1: overdispersed) for binary sampling effort and Pagel’s λ (0: phylogenetically random sampling distribution; 0-1: intermediate phylogenetic signal; 1: phylogenetic clustering under Brownian motion) for the log_10_-transformed number of samples. Finally, we applied a graph-partitioning algorithm, phylogenetic factorization, using the *phylofactor* package (99), to both the above sets of phylogenies. This approach partitioned binary sampling effort (binomial response) and the log_10_-transformed number of samples (Gaussian distribution) across the trees using iterative GLMs, producing a list of bat or rodent clades (containing at least three species) that differ significantly from the rest of the tree in sampling effort. We used Holm’s sequentially rejective test (100) with a family-wise error rate of 5% to account for multiple comparisons and determine the number of statistically significantly different clades. Trees were visualized using the *ape*, *treeio,* and *ggtree* packages (101, 102, 103).

## Acknowledgments

MMJ and DJB were supported by the National Science Foundation (DBI 2515340), and DJB was further supported by the Edward Mallinckrodt, Jr. Foundation. MMJ is also supported by a Gates Cambridge Scholarship enabled by grant OPP1144 from the Bill & Melinda Gates Foundation.

## Supporting information captions

**Supplementary Fig. 1.** Predicted values and 95% confidence intervals from a generalized linear model of binary sampling of a country for bat and rodent tissues (0: no sampling; 1: at least one bat or rodent tissue sample collected from country) across five global regions, generated with the *visreg* package in R.

**Supplementary Fig. 2.** Predicted values and 95% confidence intervals from a generalized linear model of number of bat and rodent tissue samples collected from countries across all global subregions, generated with the *visreg* package in R.

**Supplementary Fig. 3.** Predicted values and 95% confidence intervals from a generalized linear model of the binary tissue holding status of a country (0: no bat or rodent tissue samples; 1: at least one bat or rodent tissue sample) across five global regions, generated with the *visreg* package in R.

**Supplementary Fig. 4.** Predicted values and 95% confidence intervals from a generalized linear model of number of bat and rodent tissue samples housed in countries across all global subregions, generated with the *visreg* package in R.

**Supplementary Data S1.** Complete list of institutions included in our filtered dataset, including the number of bat, rodent, and total samples as well as the country code for each institution.

**Supplementary Data S2.** Complete list of institutions holding tissue samples from the ten rodent and ten bat species predicted to harbor the greatest number of undiscovered zoonoses.

## Notes

### Competing Interest Statement

The authors have declared no competing interest.

### Summary of Updates

Supplementary figures S1-S4 and data files S1-S2 have now been uploaded.

