## Supplementary Figures for "Biodiversity databases as underutilized resources for pathogen discovery: a quantitative synthesis of bat and rodent tissue collections in natural history museums"

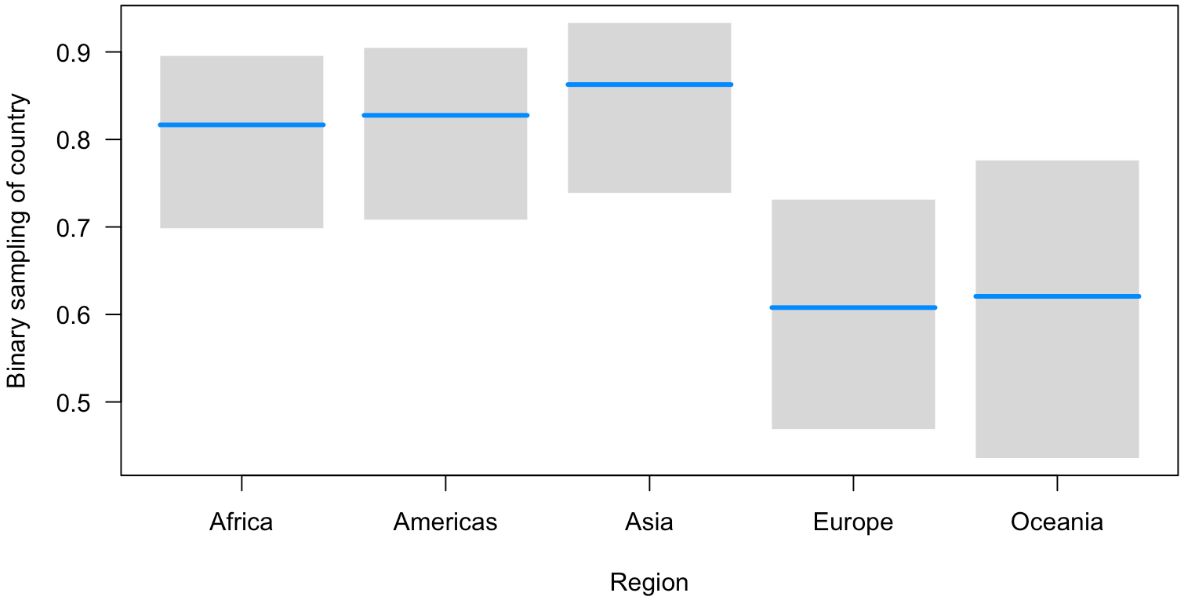


**Supplementary Fig. 1.** Predicted values and 95% confidence intervals from a generalized linear model of binary sampling of a country for bat and rodent tissues (0: no sampling; 1: at least one bat or rodent tissue sample collected from country) across five global regions, generated with the *visreg* package in R.

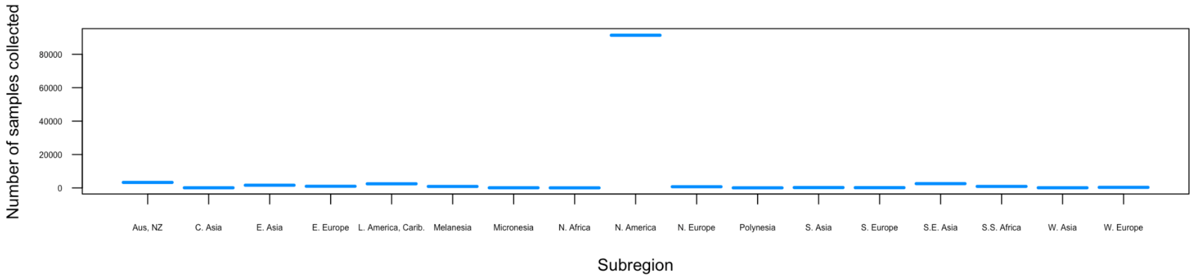


**Supplementary Fig. 2.** Predicted values and 95% confidence intervals from a generalized linear model of number of bat and rodent tissue samples collected from countries across all global subregions, generated with the *visreg* package in R.

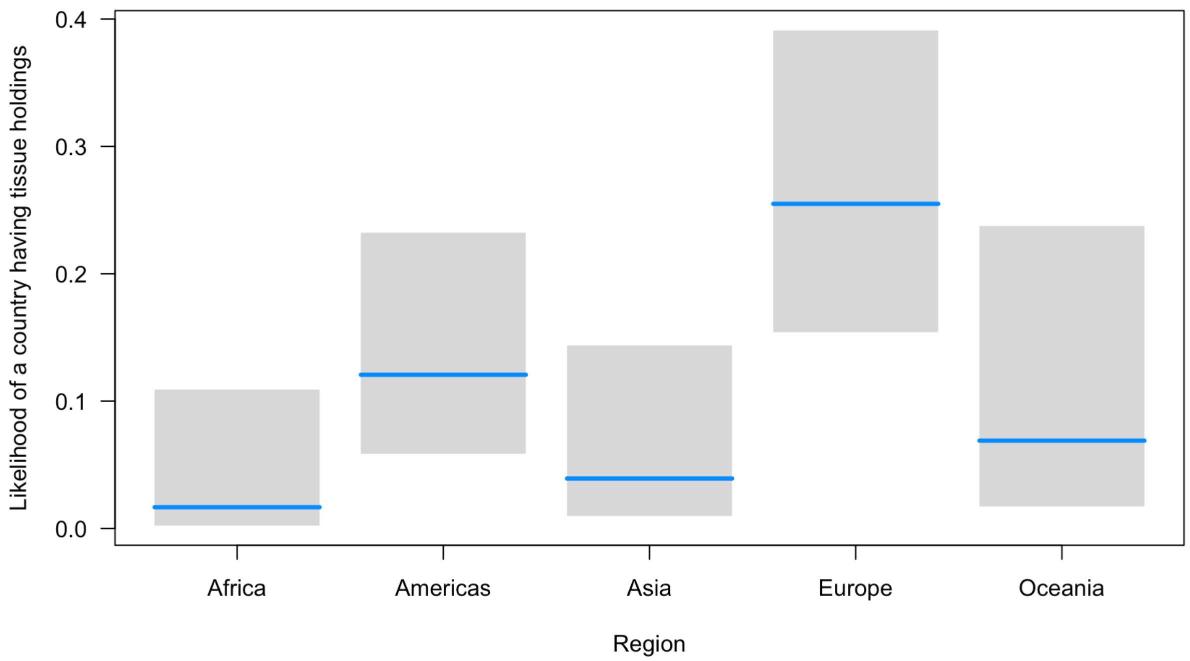


**Supplementary Fig. 3.** Predicted values and 95% confidence intervals from a generalized linear model of the binary tissue holding status of a country (0: no bat or rodent tissue samples; 1: at least one bat or rodent tissue sample) across five global regions, generated with the *visreg* package in R.

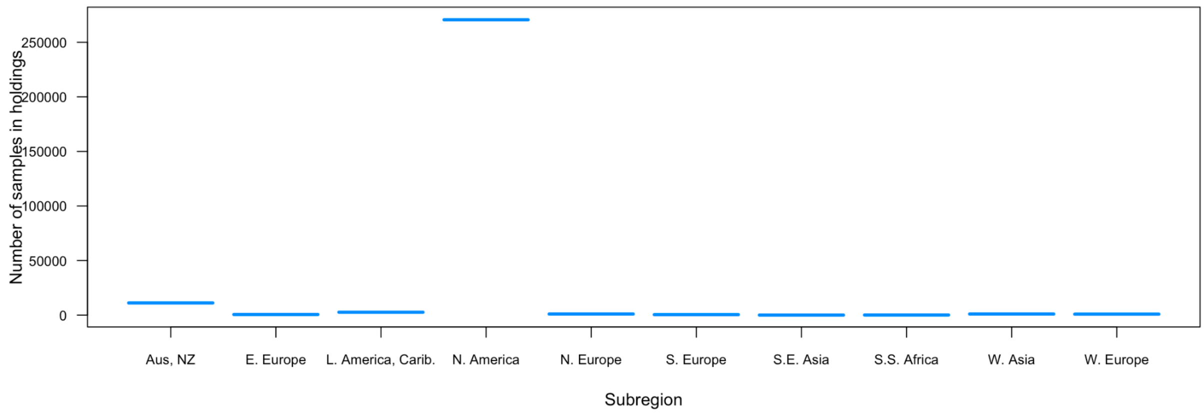


**Supplementary Fig. 4.** Predicted values and 95% confidence intervals from a generalized linear model of number of bat and rodent tissue samples housed in countries across all global subregions, generated with the *visreg* package in R.
